# N-terminal mutants of human apolipoprotein A-I: structural perturbations associated to protein misfolding

**DOI:** 10.1101/520171

**Authors:** Gisela M. Gaddi, Romina Gisonno, Silvana A. Rosú, Lucrecia M. Curto, Esteban E. Elías, Eduardo D. Prieto, Guillermo R. Schinella, Gabriela S. Finarelli, M. Fernanda Cortez, Nahuel A. Ramella, M. Alejandra Tricerri

**Affiliations:** Instituto de Investigaciones Bioquímicas de La Plata (INIBIOLP), CONICET, La Plata, Buenos Aires, Argentina; Facultad de Ciencias Médicas, Universidad Nacional de La Plata, La Plata, Buenos Aires, Argentina; Departamento de Química Biológica e Instituto de Bioquímica y Biofisica, Facultad de Farmacia y Bioquímica, Universidad de Buenos Aires, Buenos Aires, Argentina; Instituto de Investigaciones Fisicoquímicas Teóricas y Aplicadas (INIFTA) Universidad Nacional de La Plata-CONICET, La Plata, Buenos Aires, Argentina; Comisión de Investigaciones Científicas, Pcia de Buenos Aires, La Plata, Argentina; Current Address: Laboratorios de Inmunología Oncológica. Instituto de Medicina Experimental (IMEX)-CONICET-ANM, CABA, Argentina

## Abstract

Since the early description of different human apolipoprotein A-I variants associated to amyloidosis, the reason that determines its deposition inducing organ failure has been under research. To shed light into the events associated to protein aggregation, we studied the effect of the structural perturbations induced by the replacement of a Leucine in position 60 by an Arginine as it occurs in the natural amyloidogenic variant (L60R). Circular dichroism, intrinsic fluorescence measurements and assays of binding to ligands indicate that L60R is more unstable, more sensitive to proteolysis and interacts with sodium dodecyl sulfate (a model of negative lipids) more than the protein with the native sequence and other natural variant tested, involving a replacement of a Trytophan by and Arginine in the amino acid 50 (W50R). In addition, the small structural rearrangement observed under physiological pH leads to the release of tumor necrosis factor α and interleukin-1β from a model of macrophages. Our results strongly suggest that the chronic disease may be a consequence of the loss in the native conformation which alters the equilibrium among native and cytotoxic proteins conformation.

## Introduction

Human apolipoprotein A-I (apoA-I) is the main protein associated to high density lipoproteins (HDL). It is synthesized in liver and intestine and both the liver and the kidney are the major sites of its catabolism. Its multiple functions as lipid transport, endothelial homeostasis and inhibition of inflammatory pathways [1][2][3] could however be counter balanced by the presence of single point mutations which could, by not yet completely known pathways, induce its misfunction, increased clearance rate or its tendency to aggregate [4][5]. Hereditary apoA-I amyloidosis has been described since 1969, and is characterized by specific deposits of natural variant proteins within organs, in a pattern which is dependent on the mutation in the protein sequence. From the more than 20 known natural variants, almost half result from substitutions between amino acids 26 and 107, and involve with different severity hepatic and kidney failure. Instead, a “hot spot” affecting residues 173-178 was described, in which variants are especially associated to cardiac, skin and testis damage. The reasons for the different deposit patterns are still unknown but may be the consequence of long term seeding of misfolded proteins that yield a final fibrillar conformation.

The first identified apoA-I natural mutant was the result of a substitution of a Glycine by an Arginine in position 26 from the native sequence (Gly26Arg, in the abbreviated nomenclature G26R), inducing in patients peripheral neuropathy, peptic ulcer, and nephropathy [6]. The next mutations described in the N terminus of apoA-I were Trp50Arg (W50R) [7] and Leu60Arg (L60R) [8]. These variants show similarities with G26R: in each, a neutral residue is replaced by an Arginine thus increasing by one the positive net charge, and amyloid fibrils isolated from the tissues consist of the N-terminal fragments of the variant apoA-I.

As a difference from other hereditary amyloidosis, in which renal failure occurs mainly due to glomerular protein deposits [9][10], apoA-I associated disease is mostly characterized by amyloid retention in the medullary interstitium and/or vasculature, which is probably a reason for its misdiagnosis [11][12]. Only rare exceptions were observed within a few mutants including W50R, in which amyloid deposits were found in glomeruli, either confined [13], or expanded to medulla. Previous works from our and other groups have demonstrated that the single point mutations described for apoA-I decrease the marginal protein stability and elicit the tendency to aggregate; even though the conformational shift in the variants is usually subtle under physiological pH and low concentration, it could also induce alterations of the binding to ligands or the eliciting of pro-inflammatory cellular events [14][15]. In order to extend the knowledge about structural motifs that could be involved in apoA-I aggregation and pathogenicity, we herein studied and described the structural initial events that could result in high ? yield of misfolded conformations in the pro amyloidogenic variant L60R.We compared this mutant behavior with the Wt and with W50R, which is involved in renal amyloidosis but with a relative different aggregation clinical profile. This last mutant served as good comparison as some structural characterizations were already performed by Gursky and colleagues[16][17].

## Materials and methods

### Materials

Reagents purchased from Sigma-Aldrich (St Louis, MO) comprise the following: matrix metalloproteinase-12 (MMP-12, Catalytic Domain), Phorbol 12-myristate 13-acetate (TPA), Trypsin, Guanidine hydrochloride (GdmCl), sodium dodecyl sulfate (SDS), thioflavin T (ThT), polymixin B; 1,2-dimyristoylsn-glycero-3-phosphatidylcholine, (DMPC) was from Avanti Polar Lipids (Alabaster, AL). IMAC Sepharose 6 Fast Flow Resin was acquired from GE Healthcare Bio-Sciences AB, Uppsala, Sweden). The glycosamine glycan Heparin from bovine intestinal mucosa (average molecular weight 15 kDa) was from Northia (BA, Argentina). From Invitrogen (Carlsbad, CA) we obtained 4,4’-dianilino-1,1’-binaphthyl-5,5’-disulfonic acid, dipotassium salt (Bis-ANS). Isopropyl-β-D-thiogalactoside (IPTG) was purchased to Thermo Scientific (Waltham, MA). All other reagents were acquired with the highest analytical grade.

## Methods

### Cloning, expression, and purification of wild-type (Wt) W50R and L60R mutants of apoA-I

A cDNA template containing human Wt apoA-I sequence was used in order to introduce the required single point mutations. This construct was inserted into a pET-30 plasmid (Novagen, Madison, WI), allowing the expression and purification of the apoA-I variants fused to an N-terminal His-Tag peptide [18]. The mutants W50R and L60R were obtained by the Quick change method (Stratagene, La Jolla, CA). Primers were as follows:

For W50R: Sense 5′-agctccttgacaacagggacagcgtgacc-3′
Antisense 5′-ggtcacgctgtccctgttgtcaaggagct-3′
For L60R Sense 5′-caccttcagcaagcggcgcgaacagctcg-3′
Antisense 5′-cgagctgttcgcgccgcttgctgaaggtg-3

In order to further remove this N terminal peptide, an Asp—Pro sequence involving amino acid residues 2 and 3 was previously generated to allow specific chemical cleavage [19][20]. Proteins were expressed in BL 21*E Coli* strands following induction with IPTG and purified by elution through Nickel affinity columns (GE Healthcare Bio-Sciences AB, Uppsala, Sweden). The His Tag was efficiently removed by cleavage with 45% formic acid at 45^°^C for 5 h, following exhaustive dialysis against buffer Tris 20 mM, NaCl 150 mM, pH 8.0). A second metal affinity chromatography step was run to separate the final pure protein fraction.

### Mass Spectrometry Analysis

Protein digestion and Mass Spectrometry analysis were performed at the Proteomics Core Facility CEQUIBIEM, at the University of Buenos Aires/CONICET (National Research Council) as follows: Protein samples were reduced with dithiothreitol (DTT) and alkylated with iodoacetamide in Ammonium Bicarbonate 50 mM pH 8.0. This protein solution was precipitated with trichloroacetic acid (TCA). Proteins were resuspended in the same buffer, digested with trypsin (Promega V5111), peptides purified and desalted with ZipTip C18 columns (Millipore). The digests were analyzed by nanoLC-MS/MS in a Thermo Scientific Q-Exactive Mass Spectrometer coupled to a nanoHPLC EASY-nLC 1000 (Thermo Scientific). For the LC-MS/MS analysis, approximately 1 μg of peptides was loaded onto a reverse phase column (C18, 2 μm, 100 A, 50 μm x 150 mm) Easy-Spray Column PepMap RSLC (P/N ES801) suitable for separating protein complexes with a high degree of resolution. The MS equipment has a high collision dissociation cell (HCD) for fragmentation and an Orbitrapanalyzer (Thermo Scientific, Q-Exactive). XCalibur 3.0.63 (Thermo Scientific) software was used for data acquisition and equipment configuration that allows peptide identification at the same time of their chromatographic separation. Full-scan mass spectra were acquired in the Orbitrap analyzer. Q-Exactive raw data were processed using Proteome Discoverer software (version 2.1.1.21 Thermo Scientific) and searched against apoA-I sequence database downloaded from NCBI (National Center for Biotechnology Information) (www.ncbi.nlm.nih.gov) digested with trypsin with a maximum of one missed cleavage per peptide.

The exponentially modified protein abundance index (emPAI) was calculated automatically by Proteome Discoverer software and used to estimate the relative abundance of identified proteins within the sample.

### Protein structure and chemical stability

ApoA-I Wt, W50R and L60R were diluted in Tris 20 mM pH 7.4 (Tris buffer) or citrate phosphate McIlvaine’s buffer(Citrate buffer) pH 5.0 [21]. Variants were taken at 0.2 mg/mL (at 25°C). Tryptophan (Trp) intrinsic fluorescence emission spectra was acquired in an SLM4800 spectrofluorometer (ISS Inc, Champaign, IL) upgraded by Olis, setting the excitation wavelength at 295 nm and scanning emission from 310 to 400 nm. Solvent exposure of Trp residues was determined as a parameter of protein structural arrangement. ApoA-I variants were diluted to 0.1 mg/mL (in Tris buffer). Quenching of the Trp residues was determined by measuring intrinsic fluorescence following stepwise addition of acrylamide from a concentrated stock solution. The quenching constant K was calculated from a linear plot of the Stern-Volmer equation as:

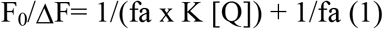

where fa is the fraction of the initial fluorescence which is accessible to the quencher, K is the Stern-Volmer quenching constant of the accessible fraction and [Q] is the concentration of the quencher. F_0_ is the initial fluorescence in the absence of quencher and ΔF is the remaining fluorescence after the addition of acrylamide at each concentration [22][15].

In order to sense hydrophobic pockets within the proteins spatial arrangements, the binding of Bis-ANS was measured following titration of the probe to apoA-I variants. Proteins were set at 0.1 mg/mL in Tris buffer and Bis-ANS added in small amounts from a methanol stock. Both, total intensity and spectral shift were registered by excitation at 395 nm and emission scanned from 450 to 550 nm [15]. Circular dichroism (CD) measurements were acquired on a Jasco J-810 spectropolarimeter. To register far UV spectra (200–290 nm), proteins were diluted in phosphate 10 mM buffer, pH 7.4 to 0.2 mg/mL in a 1 mm path-length cuvette, and three scans at a speed of 20 nm min^-1^ were registered and averaged to minimize noise signal. Mean residue weight values of 115.5, 115.4 and 115.7 for Wt, were estimated to calculate molar ellipticity as previously described [23]. Near CD spectra were registered under the same conditions but with a final protein concentration of 1.0, 1.5 and 1.5 mg/mL for Wt, W50R, and L60R, respectively.

To characterize apoA-I variants’ stability respect to the Wt, Trp fluorescence spectra were measured following stepwise additions of increasing amounts of GdmCl [15][14]. As described in our previous works, the free energy of unfolding in the absence of denaturant (ΔG°H2O) was calculated from the shift of the Trp spectral center of mass as a function of GdmCl final concentration assuming the simplest two state process model [18][19].

In order to compare the self-association of the proteins in solution, Wt, W50R, and L60R were taken to 0.5 mg/mL in Phosphate Buffer (pH 7.4) and eluted through a Sepharose 200 HR column (Amersham Pharmacia recommended MW 10,000-600,000), at a Flow rate of 0.35 mL/min, connected to a Merck Hitachi Fast Performance Liquid Chromatography equipment. Elution profile was followed by UV detector (280 nm).

### Partial degradation by proteolysis

In order to compare the accessibility of the variants to partial proteolysis, proteins were incubated at 37^°^C in Tris buffer with Trypsin at a molar ratio apoA-I variants to enzyme 1000:1. At different periods samples were heated in boiling water, resolved by SDS PAGE, and developed by silver staining. The associated intensity of the protein remaining within the monomer molecular weight was quantified with the Image J 1.51 j8 Software. In the same trend, proteins were incubated with metalloproteinase 12 (molar ratio protein to enzyme 500:1) and analyzed in the same way.

### Detection of protein aggregates

Thioflavin T (ThT) is widely used to detect amyloid-like structures [24]. Although ThT-associated quantum yield usually increases proportionally to the formation of fibrillar protein aggregates, significant fluorescence is detected even when proteins are present as oligomeric conformations [25][26]. To address the effect of pH on the yield of amyloid-like protein aggregates, Wt (0.2 mg/mL) was taken for 48 h at 37°C at different pH between 7.4 and 5.0 in Citrate phosphate buffer, and ThT added at a molar ratio of two with respect to protein. Associated fluorescence intensity was measured on a Beckman Coulter DTX 880 Microplate Reader (Beckman, CA) through the excitation (430 nm) and emission (480 nm) filters. The relative size of the aggregates was estimated by light scattering on theSLM4800 spectrofluorometer measuring light intensity at 90^°^ with excitation and emission wavelength set at 400 nm. Next, to compare the efficiency of the variants to bind to heparin, 0.2 mg/mL of Wt, W50R, and L60R, were incubated for 48 h at 37°C, pH 5.0. Both fluorescence and scattering were determined in the Microplate Reader, in the latter case by fixing excitation and emission filters at 350 nm.

To characterize nanometer-scaled aggregates conformations, Transmission electron microscopy (TEM), was performed on a JEOL-1200 EX. Negative staining is widely used to increase the contrast as the stain surrounds the sample but is excluded from the occupied volume and is thus observed as ‘negative stain’ [27]. Samples incubated at 0.6 mg/mL (37°C for 7 days) were seeded on Formvard grids, 0.5% phosphotungstic acid was added and visualized by negative staining. Magnification was 100,000 x.

## Lipid interaction properties of apoA-I variants

Sodium dodecyl sulfate (SDS) is a negative lipid which mimics some characteristics of biological membranes [28]; in addition, it has been described to elicit the formation of fibrils from different peptides and proteins if used below the critical micellar concentration (CMC). We have previously shown that Wt binds to this lipid increasing ThT fluorescence [15]. To analyze the comparative binding of SDS to apoA-I variants, proteins were incubated in citrate phosphate buffer pH 7.4 for 48 h at 37°C in the presence or absence of 0.2 mM SDS. Previously we have confirmed that under those conditions SDS is far lower from the CMC. ThT-associated fluorescence and scattering were measured as described above.

DMPC clearance assay was used to give account for apoA-I function in lipid solubilization. As proteins are incubated with this phospholipid at its transition temperature the decrease in turbidity indicates the efficiency to form lipid-protein complexes. DMPC was solubilized in chloroform and desired amounts were dried under extensive N_2_ flow and additional vacuum. Lipids were resuspended in Tris buffer and Multilamellar liposomes (MLV) were obtained by exhaustive vortexing. Lipid clearance was determined by incubating proteins with DMPC MLV at a molar ratio lipid:protein 40:1 at 24°C for 90 min and absorbance monitored on the Microplate Reader by setting filters at 350 nm [15].

## Pro-inflammatory response induced by variants

In order to determine whether a soluble conformation of W50R and L60R could elicit cellular pro-inflammatory pathways, the human-derived THP-1 Cell Line (from Leukemic monocytes, ECACC, Salisbury, UK) was seeded (10^6^ cells/mL) in RPMI 1640 medium in the presence of 10% fetal bovine serum (FBS), 100 U/mL Penicillin, and 100 ug/mL Streptomycin at 37°C in a humidified incubator containing 5% CO_2_. Monocytes were activated by the addition of 5 ng/mL of Phorbol esters for 48 h [29]. Transformation into macrophages was observed by cellular adhesion to the plate. Thereafter, medium was removed and apoA-I variants (at 1.0 μg/mL) added to the cells and incubated for3 h in RPMI medium plus 0.5% FBS, in the presence of the same antibiotic mixture and Polymyxin B (final concentration 50 μg/mL). Positive and negative controls were determined by addition of 50ng/mL of bacterial lipo polysaccharide (LPS) in the absence of the presence of Polymyxin B, respectively. Cells were spun at 2,100 rpm for10 min to remove cellular debris, supernatant separated and tumor necrosis factor α (TNF-α) and interleukin-1β (IL-1β) release were compared by specific enzyme immunoassay from BD Biosciences (San Diego, CA) used according to the manufacturer’s instructions. Cell viability under these incubation conditions was checked by the 3-(4,5-dimethylthiazolyl)-2,5-diphenyl-tetrazoliumbromide (MTT) cell viability assay as previously described [14].

## Other analytical methods

Protein content was quantified by the Bradford technique [30] or by absorbance from the estimation of the extinction coefficient (32,430 M^-1^cm^-1^at 280 nm) as determined in a Bio-Rad spectrophotometer (Hercules, CA).Unless otherwise stated, the results were reproduced in three independent experiments and are indicated as means ± standard error. Statistically significant differences between experimental conditions were evaluated by the Student’s test.

## Results

### Structural comparison and stability

The expression and isolation procedures used in this study yielded high amounts of pure proteins. We have previously shown that the lack of the two first amino acids resulting from the acidic cleavage does not introduce modifications in protein function or structure with respect to the plasma apoA-I [20]. In order to confirm the chemical integrity and purity, the Wt variant was subjected to LC-MS/MS equipped with an Orbitrap analyzer. Exponentially Modified Protein Abundance Index (emPAI) estimated an abundance of 99.95 for this protein. Previous to each experiment, in order to ensure a fresh folding, proteins were solubilized in GdmCl 2 M and extensively dialyzed through the desired buffer.

Due to the location in the primary structure of the four Trp residues in apoA-I molecule (8, 50, 72 and 108), the average intrinsic fluorescence is representative of the conformation of the N-terminal domain (residues 1-184) [31]. In order to compare proteins structure, we prepared freshly refolded apoA-I variants and characterized Trp fluorescence spectra. While the wavelength of maximum fluorescence (WMF) of W50R was similar to the Wt, a small but significant red shift (4 nm +/-1) was observed for L60R (Table 1).

**Table 1.**
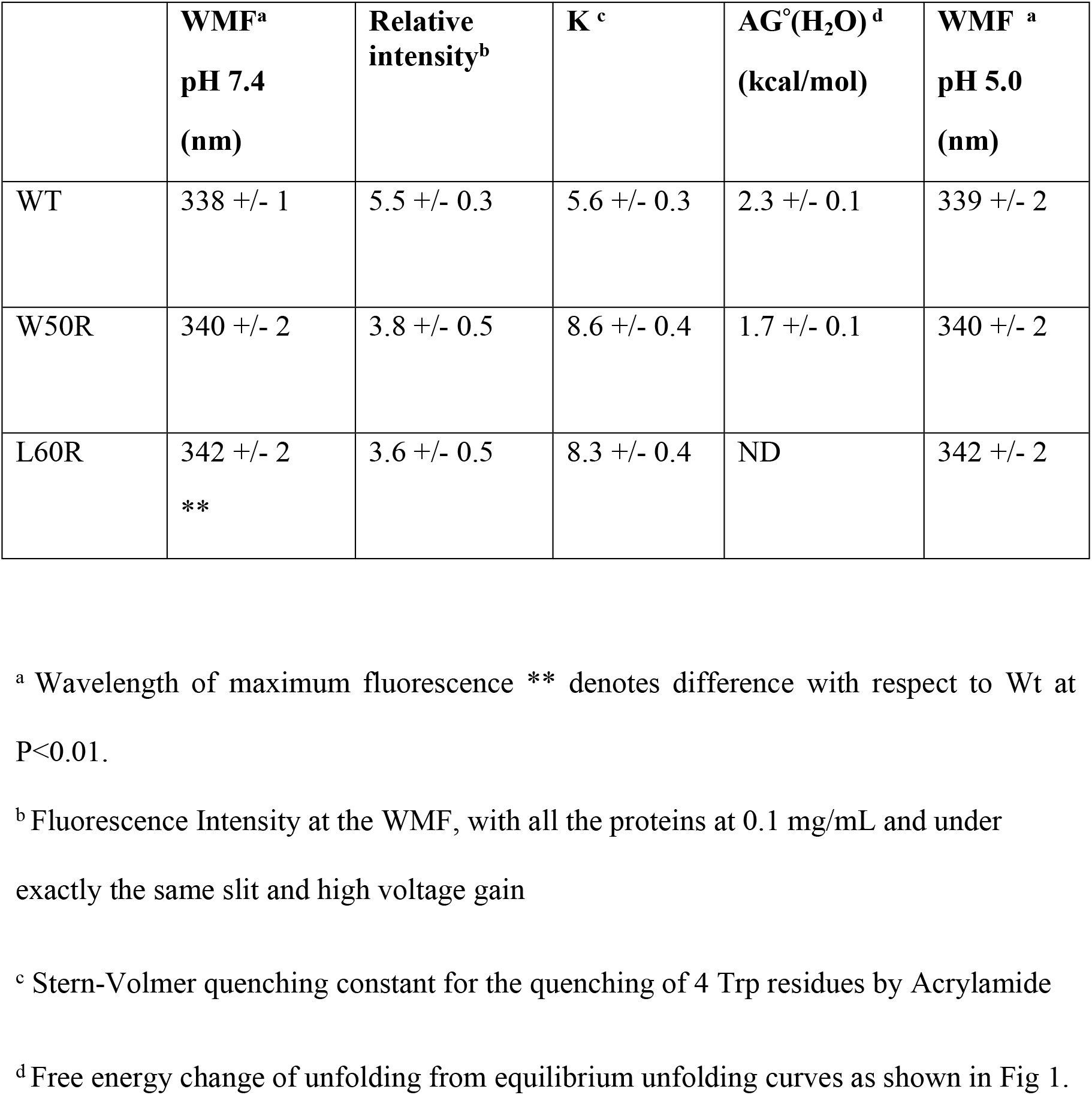
Spectral characterization of N terminal apoA-I variants.

|  | **WMF<sup>a</sup>**<br>**pH 7.4**<br>**(nm)** | **Relative**<br>**intensity<sup>b</sup>** | **K<sup>c</sup>** | **AG<sup>o</sup>(H<sub>2</sub>O)<sup>d</sup>**<br>**(kcal/mol)** | **WMF<sup>a</sup>**<br>**pH 5.0**<br>**(nm)** |
| --- | --- | --- | --- | --- | --- |
| WT | 338 +/- 1 | 5.5 +/- 0.3 | 5.6 +/- 0.3 | 2.3 +/- 0.1 | 339 +/- 2 |
| W50R | 340 +/- 2 | 3.8 +/- 0.5 | 8.6 +/- 0.4 | 1.7 +/- 0.1 | 340 +/- 2 |
| L60R | 342 +/- 2<br>** | 3.6 +/- 0.5 | 8.3 +/- 0.4 | ND | 342 +/- 2 |

<sup>a</sup> Wavelength of maximum fluorescence \*\* denotes difference with respect to Wt at P<0.01.
<sup>b</sup> Fluorescence Intensity at the WMF, with all the proteins at 0.1 mg/mL and under exactly the same slit and high voltage gain
<sup>c</sup> Stern-Volmer quenching constant for the quenching of 4 Trp residues by Acrylamide
<sup>d</sup> Free energy change of unfolding from equilibrium unfolding curves as shown in Fig 1.

Both mutants showed lower intensity than the Wt for identical protein concentration. A significantly higher Stern-Volmer constant (K) calculated from the analysis of Trp quenching with acrylamide (8.6+/- 0.4 for W50R and 8.3+/- 0.4 for L60R) indicates a higher exposure of the Trp environments to solvent in both mutants as compared to the Wt form (K 5.6 +/- 0.3).

To better estimate apoA-I natural variants’ conformational stability, fluorescence was followed after titration with GdmCl. The chemical denaturation pattern obtained from the shift in the center of mass of the Trp emission has been extensively used to give account for the protein stability under chaotropic solvents [32]. As previously discussed, this parameter is in good agreement with the measurement of the emission intensity at a fixed wavelength at increasing GdmCl concentration, and better help to compare our previous data [14][15]. While the free energy of denaturation (ΔG°H_2_O) for W50R (1.7 kcal/mol) is lower than the value estimated for the Wt (2.3 kcal/mol), the dependence fit of Trp emission with GdmCl is far from a two state (native and unfolded) model for L60R (Fig 1A), and thus we were not able to precisely calculate this parameter. Interestingly, its total intrinsic fluorescence intensity is lower than the one of the Wt, even though it keeps the four native Trp residues. As long as protein is unfolded by titration with GmdCl the total intensity tends to equal that of the Wt (Inset Fig 1A).

**Fig 1.**
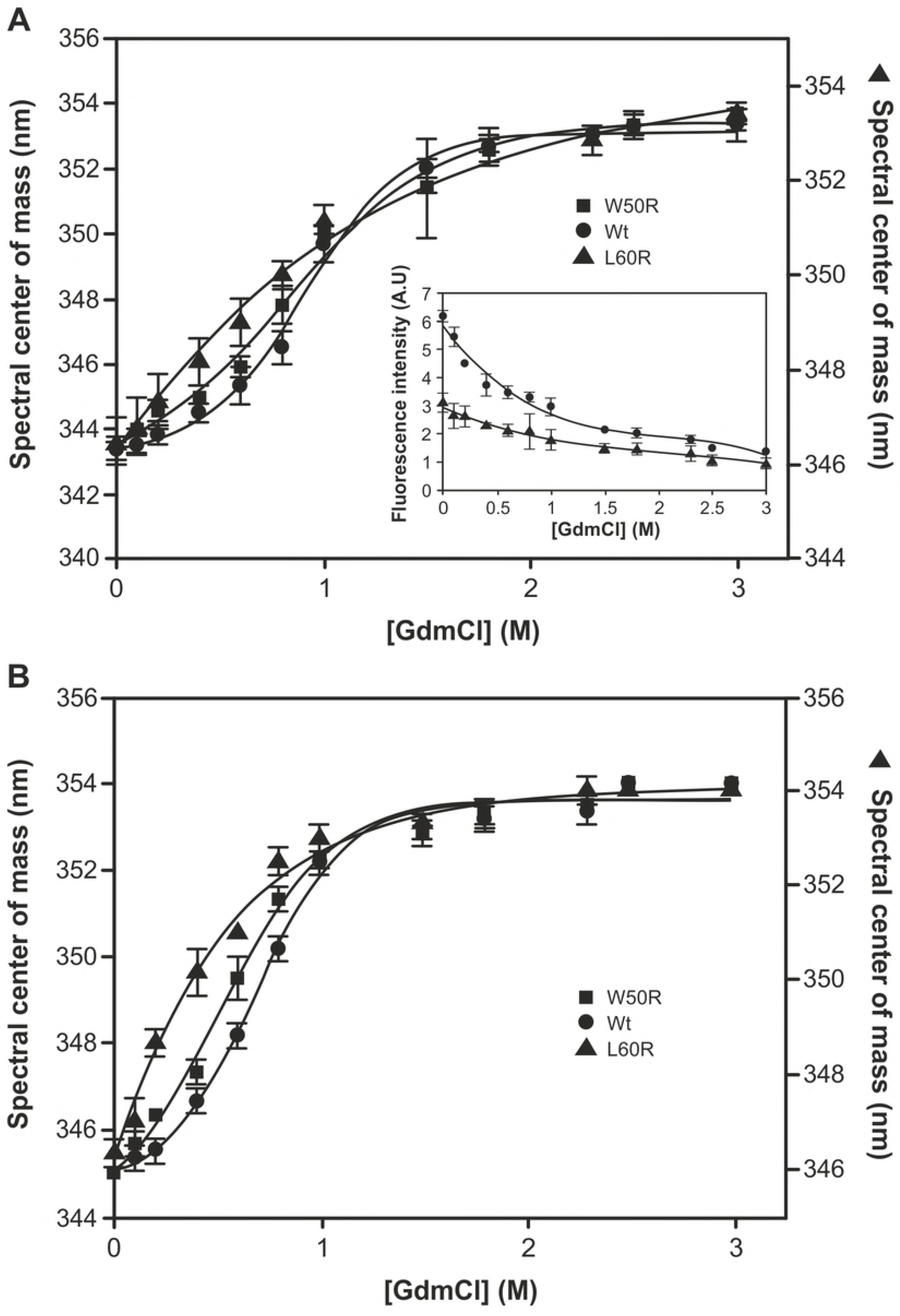
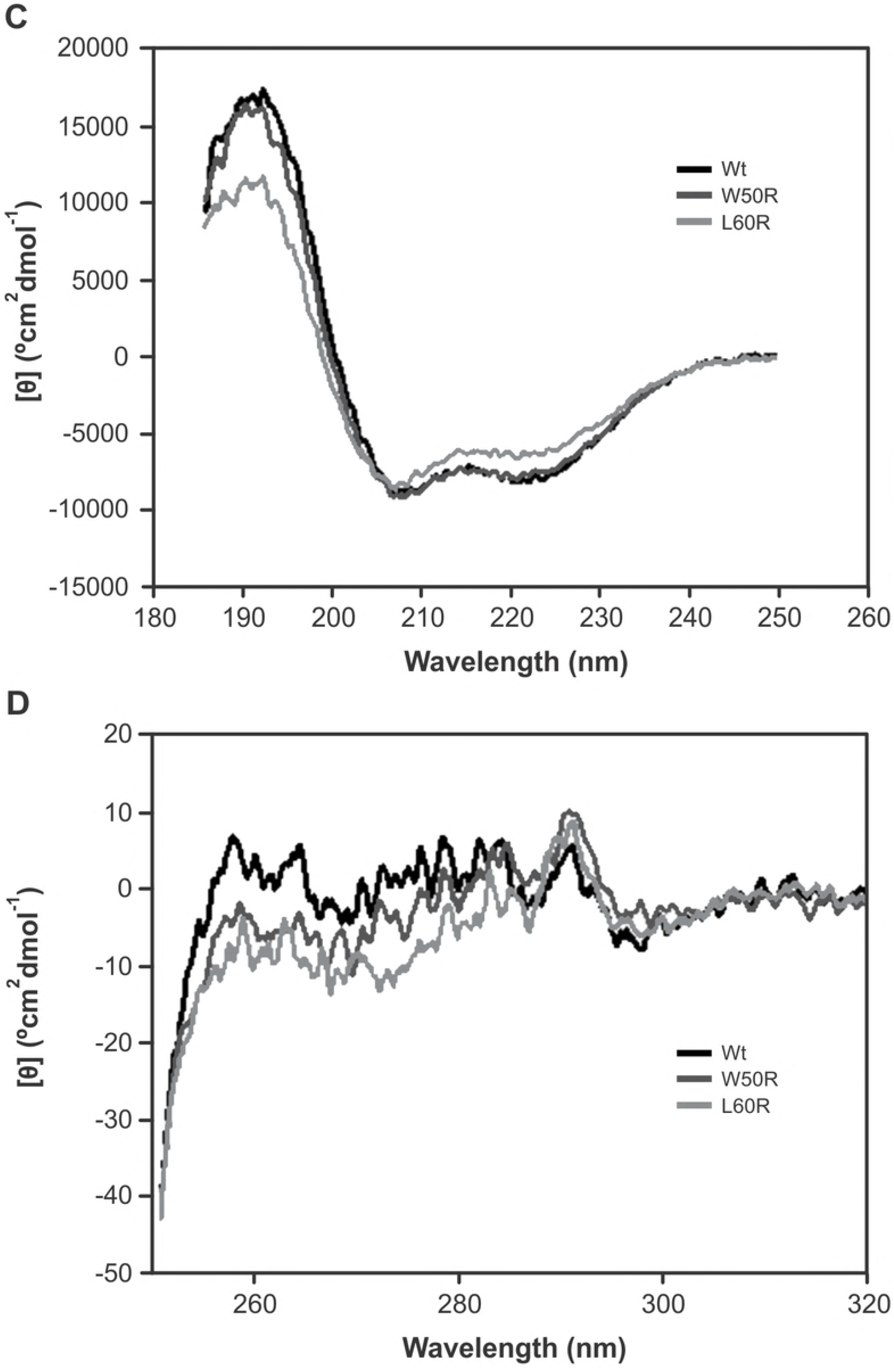
Structural characterization of apoA-I variants. Chemical equilibrium unfolding of apoA-I in Tris 20 mM buffer pH 7.4 (A) and pH 5.0 in Citrate phosphate buffer (B) was evaluated at a starting protein concentration of 0.2 mg/mL; Trp fluorescence was registered with excitation at 295 nm and emission between 310 and 400 nm; spectral center of mass was plotted as a function of [GmdCl]. Circles, squares and triangles represent Wt, W50R, and L60R, respectively. Lines correspond to fittings of a sigmoideal model to the data. Y-left axe corresponds to Wt and W50R, and Y-right axe to L60R. Inset Fig A represents the dependence of the total intensity of L60R respect of Wt as a function of [GmdCl]. Circular dichroism in the far (C) or near (D) UV region are represented for Wt (dark), W50R (dark grey) and L60R (clear grey).

In order to test whether a local decrease in pH could induce a disruption in protein structure, variants were taken to pH 5.0 and denaturation curves repeated under the same conditions. As Fig 1B shows, the behavior remains similar to that obtained under physiological pH, indicating a mild effect of an acidic milieu on proteins’ conformation.

CD analysis of protein structure in the far UV reveals a conserved α helical secondary structure which was previously identified for the Wt protein at low concentrations (Fig 1C) [33]. The secondary structure content was calculated with the algorithm CONTIN rendering a high percentage of helical structure for all the three proteins [34]. The Wt spectrum is almost indistinguishable from that of W50R and slightly more intense than that of L60R. Accordingly, a loss of about 4 and 7% of alpha helical structure is observed for the W50R and L60R variants, respectively.

The comparison among the tertiary structures was analyzed by CD in the near UV (Fig 1D); spectra of both mutants preserve fine structure although the lower intensity from 280 nm to 250 nm indicates mild structural differences among the aromatic amino acid residues.

Partial proteolysis sensitivity is usually evaluated in order to give information about the accessibility of protein domains within the spatial arrangement. Thus, variants were incubated at increasing times with either trypsin or metalloproteinase 12 (MMP12), and run through an SDS PAGE visualized with Silver staining; the time dependent efficiency of the proteolysis could be observed and quantified by the disappearance of the band corresponding to the original Molecular weight (Fig 2A). The digestion rate of the major fragment is higher as long as the incubation time increases indicating the efficiency of the proteolysis.

**Fig 2.**
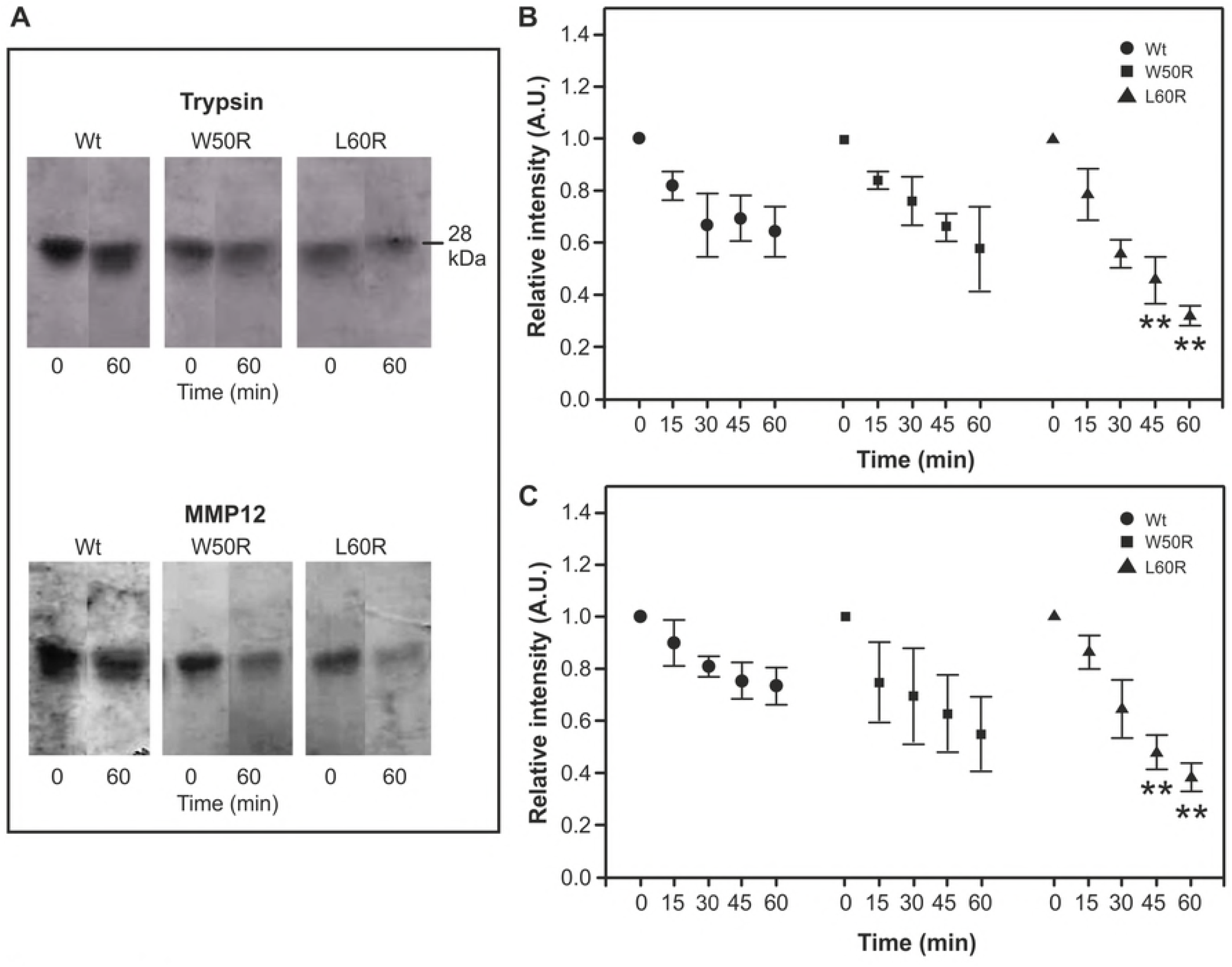
Partial proteolysis of apoA-I variants. Proteins were incubated at 0.3 mg/mL in Tris 20 mM buffer pH 7.4 with trypsin or MMP12 at molar ratios of 1000:1 or 500:1apoA-I variants to enzyme respectively. After different time periods, reactions were stopped by the addition of sample running buffer and two minutes boiling. Samples were run through a SDS PAGE (16%) and developed by silver staining. A) Black/white representation of the initial (0) and final (60 min) incubation times. Intensity remaining with the monomeric molecular weight (28 kDa) after B) Trypsin or C) MMP12 treatments was quantified by the Image Quant software and normalized to the intensity of the band at time= 0 for each protein. Circles, squares and triangles correspond to Wt, W50R and L60R respectively (** represents P< 0.001 with respect to the same time in the Wt).

After 45 minutes incubation under these conditions and in agreement with Das et al [16] the proteolysis of W50R showed a similar yield as the Wt, instead it demonstrated to be significantly more efficient for L60R with both enzymes tested.

The extrinsic fluorescence probe ANS (or its dimer Bis-ANS) was extensively used to test the spatial arrangements of proteins, due to the fact that is it weakly fluorescent in water but its quantum yield increases significantly, and its emission shifts upon binding to proteins. Its emission is proposed to sense specifically protein hydrophobic pockets and “molten globule” like states [32]. We and others have previously shown that binding of Bis-ANS to apoA-I Wt is efficient and fast [14][15]. We herein show (Fig 3), as it was previously demonstrated with the monomer ANS, that W50R binds Bis-ANS with a relatively higher quantum yield than the Wt [17].

**Fig 3.**
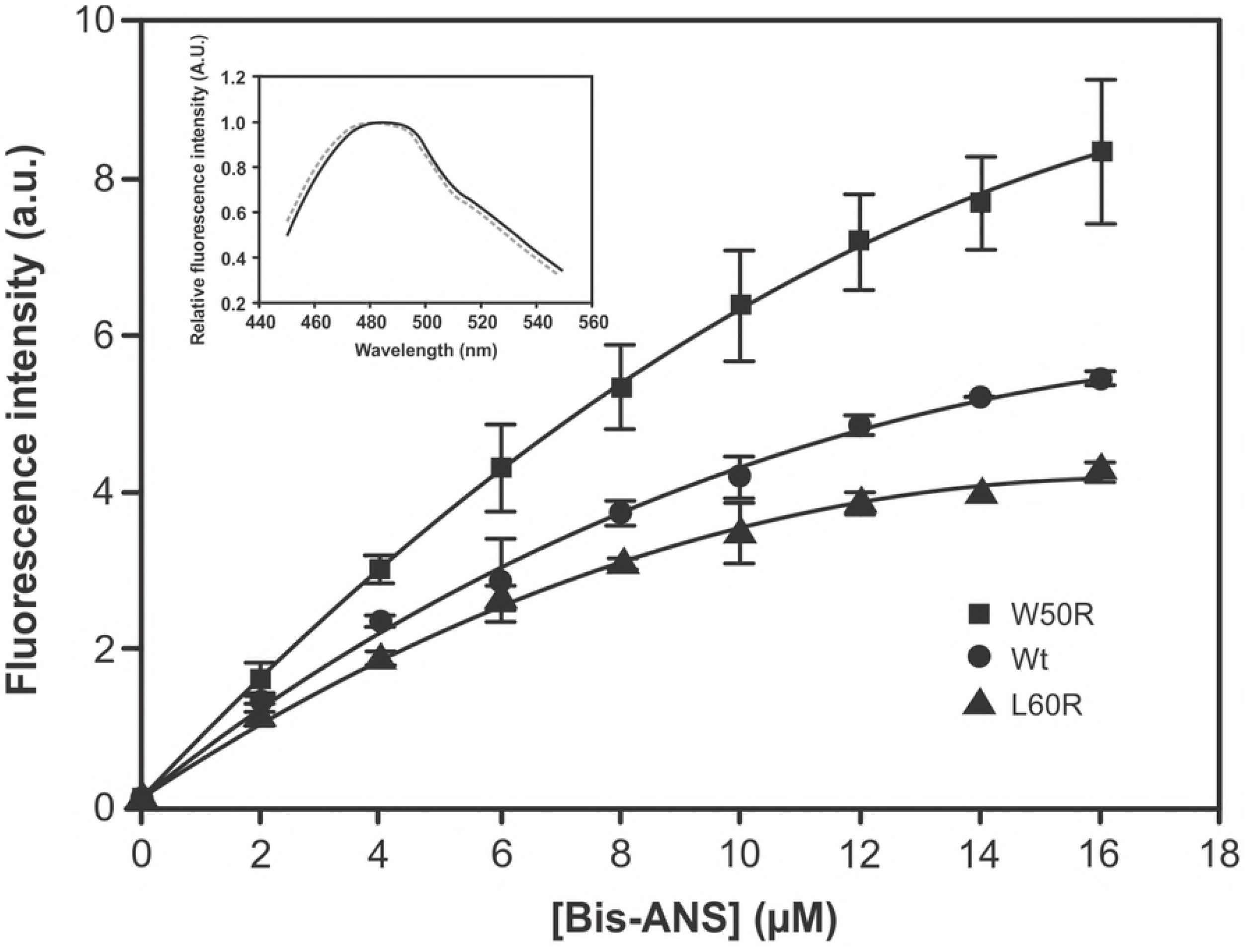
Binding of Bis-ANS to apoA-I variants. Proteins were diluted at a final concentration of 0.1 mg/mL (Tris 20 mM pH 7.4 buffer) and titrated with small amounts of Bis-ANS to a final concentration of 16 μM. Fluorescence was detected in the SLM 4800 spectrofluorometer setting excitation at 360 nm, and intensity of the emission registered at the observed Wavelength of Maximum Fluorescence (WMF) for the probe (490 nm). As in Fig. 1 circles, squares and triangles represent Wt, W50R, and L60R, respectively. Inset: spectra of Bis-ANS at a molar ratio 4:1 probe to protein normalized at the WMF. Continuous and dashed lines represent Wt and L60R respectively.

Instead, intensity of the Bis-ANS associated to L60R is lower indicating a lower permeation or binding of this probe within the spatial protein arrangement. Nevertheless, WMF of the Bis-ANS bound to L60R is similar to the probe associated to the Wt (Fig 3 inset). In a different experiment, to evaluate whether the presence of the hydrophilic group in the mutants may disrupt the protein self-association, freshly folded Wt and variants were run under FLPC; the elution pattern demonstrated that most of the Wt migrates as a dimer with a minor fraction (about 20%) eluting as a monomer (S. Fig 1). This pattern is well preserved for both mutants.

### Aggregation tendency

The variants tendency to aggregate under physiological pH and low concentrations was evaluated by incubating proteins at 0.2 mg/mL for 48 h in Citrate buffer pH 7.4. The ThT measurement indicates as previously observed for the Wt that amyloid complexes were not detected under these conditions (dark gray bars in S. Fig 2). In a parallel experiment, proteins were further incubated at 0.6 mg/mL for 7 days under the same conditions and observed by Transmission Electron Microscopy (Fig 4).

**Fig 4.**
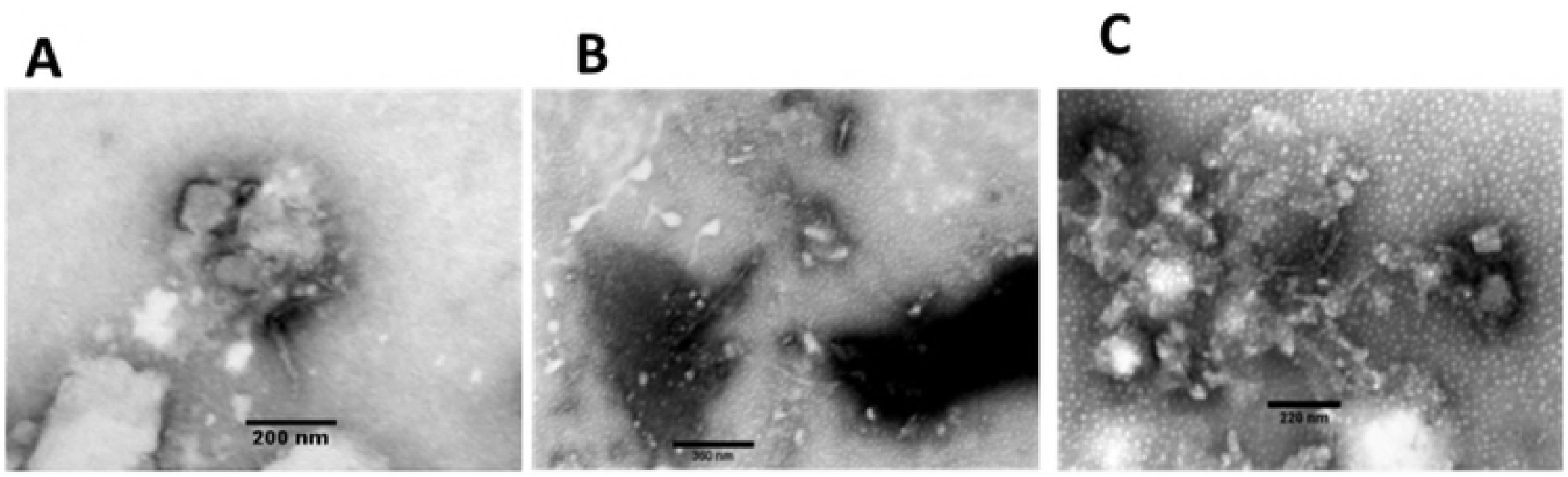
Observation of apoA-I mutants’ aggregates at physiological pH. Wt (A), W50R (B) or L60R (C) were incubated for 7 days at 0.6 mg/mL and 37 °C in Citrate phosphate buffer pH 7.4. Aggregates were observed by negative staining under Transmission Electron Microscopy on a JEOL-1200 EX Microscope. Bars in each image show the scale used.

The sample of the Wt showed a low degree of amorphous-like aggregates and small amounts of protofibers (Fig 4A). Both mutants (W50R and L60R, Fig 4B and C respectively) were represented by a higher yield of aggregates with similar morphology. In addition protofibers of about 8-10 nm diameter are observed under these conditions. The behavior of these proteins was similar to other mutants analyzed under the same conditions [14][15].

### Binding to ligands

Heparin is usually used as a model of glycosamino glycans (GAGs), which are supposed to play key roles in the maintenance of cellular functions. Moreover, it is proposed that interactions of proteins with GAGs could result in the retention or aggregation of proteins inducing amyloidosis [35][36], but instead they could compete avoiding cellular toxicity or conformational shifts [37]. Within this frame in mind, we analyzed binding of both natural mutants to heparin either at physiological pH or under acidic conditions. ThT results suggest, as observed before for the Wt under these conditions [19], that the presence of heparin does not induce a significant structural arrangement of the variants when incubated at pH 7.4 for 2 days at 37^°^C (clear gray bars in S Fig 2). As long as pH decreases, protonation of His residues may result in the gain of positive charges of the proteins thus helping electrostatic interactions with the negative groups of the GAGs. As the isoelectric point of apoA-I is estimated in 5.27, we analyzed the effect of an acidic environment on apoA-I aggregation and interaction with heparin by incubating Wt at different pH between 5.0 and 7.4, in the absence or the presence of this model of GAG. Size and amyloid-like tendency of the complexes were followed by scattering (Fig 5A) and ThT fluorescence (Fig 5B). These data indicate that the formation of Wt amyloid-like complexes is favored as long as pH decreases (dark gray bars), and this tendency is increased in the presence of heparin (clear gray bars).

**Fig 5.**
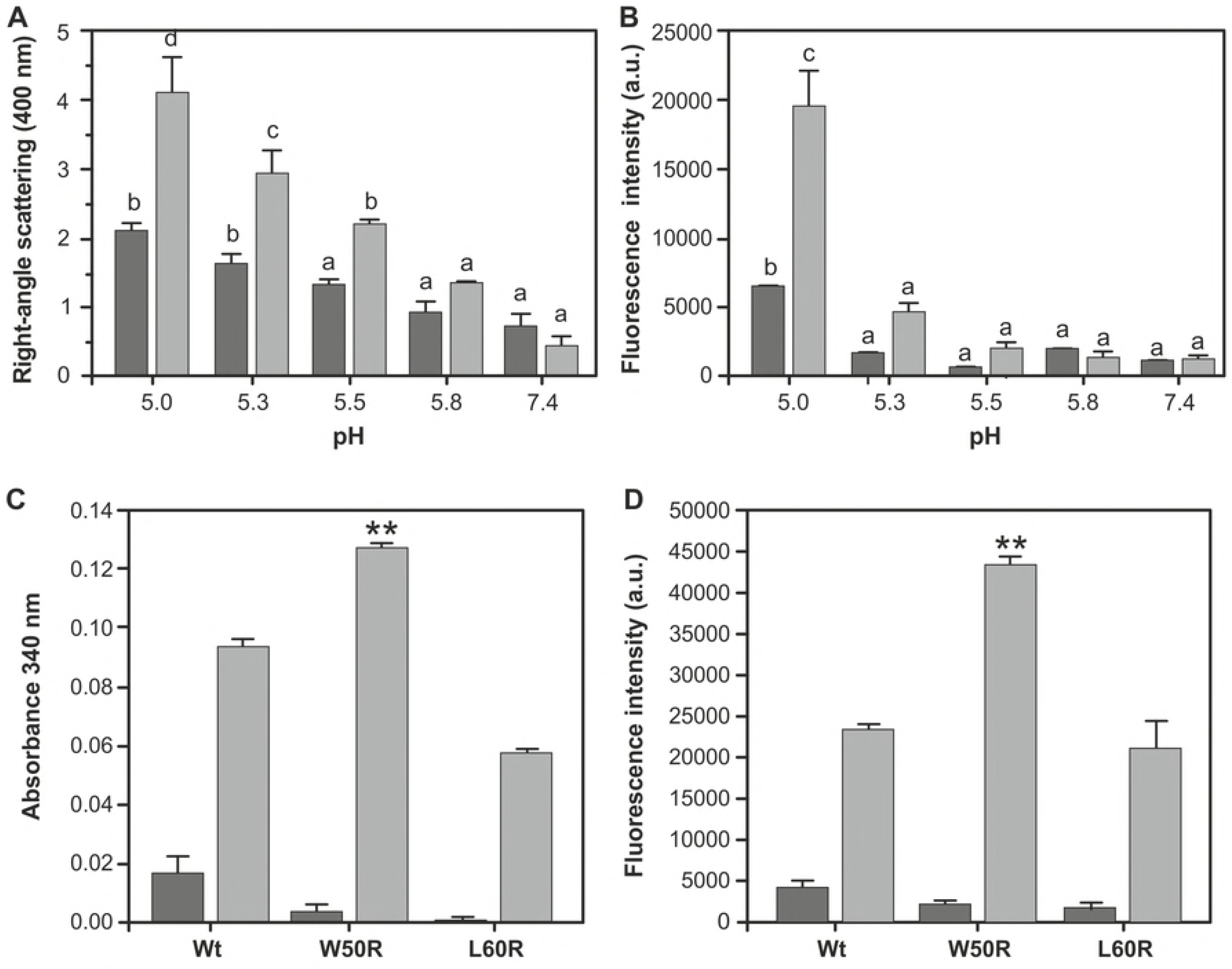
Influence of acidic pH on heparin binding. A and B) Wt was solubilized at 0.2 mg/mL at different pH from 7.4 to 5.0 by using Citrate phosphate buffer and incubated either in the absence (dark gray bars) or the presence (clear gray bars) of heparin at a molar ratio 1:1 protein to GAG for 48 h at 37°C. A) Light scattering at 90^°^ was determined in the spectrofluorometer with excitation and emission wavelengths at 400 nm; B) ThT associated fluorescence was determined in the Multiplate Reader with excitation set at 430 and emission filter at 480 nm. Bars correspond to mean ± SD. Differences were analyzed by ANOVA followed by Tukey test. Different letters symbolize significant differences (P<0.05). C and D) Wt, W50R, and L60R, were taken at pH 5.0 in Citrate phosphate buffer (0.2 mg/mL) and incubated in the absence (dark gray bars) or the presence (clear gray bars) of heparin at a molar ratio of 2 per mol of protein at 37°C by 48 h. C) Scattering was analyzed in the Microplate Reader with filter set a 340 nm; D) ThT associated fluorescence was quantified as described in Fig 5B. Symbol ** in C and D corresponds to differences with P<0.001 with respect to Wt under the same conditions.

Next, we compared the binding of the variants under study to this model of GAG at pH 5.0. As shown, a higher scattering (Fig 5C) and ThT associated fluorescence (Fig 5D) in the case of W50R indicates its higher efficiency to bind to heparin as amyloidlike complexes under acid conditions while L60R behaves similar to Wt.

In order to answer whether the structural disruption induced by a change in the amino acid sequence (and charge) of the apoA-I variants could modify their interactions with negative ligands in the microenvironment, we incubated W50R and L60R with SDS, which not only is a good model of membrane lipids but also was suggested to work as inductor of amyloid-like complexes formation from different proteins when incubated under the lipid critical micellar concentration (CMC: 0.7 mM) [38]. Fig 6 shows, as previously observed, a strong increase in ThT fluorescence of Wt protein when bound to SDS [15] which is in a similar trend for W50R and with a higher yield for L60R.

**Fig 6.**
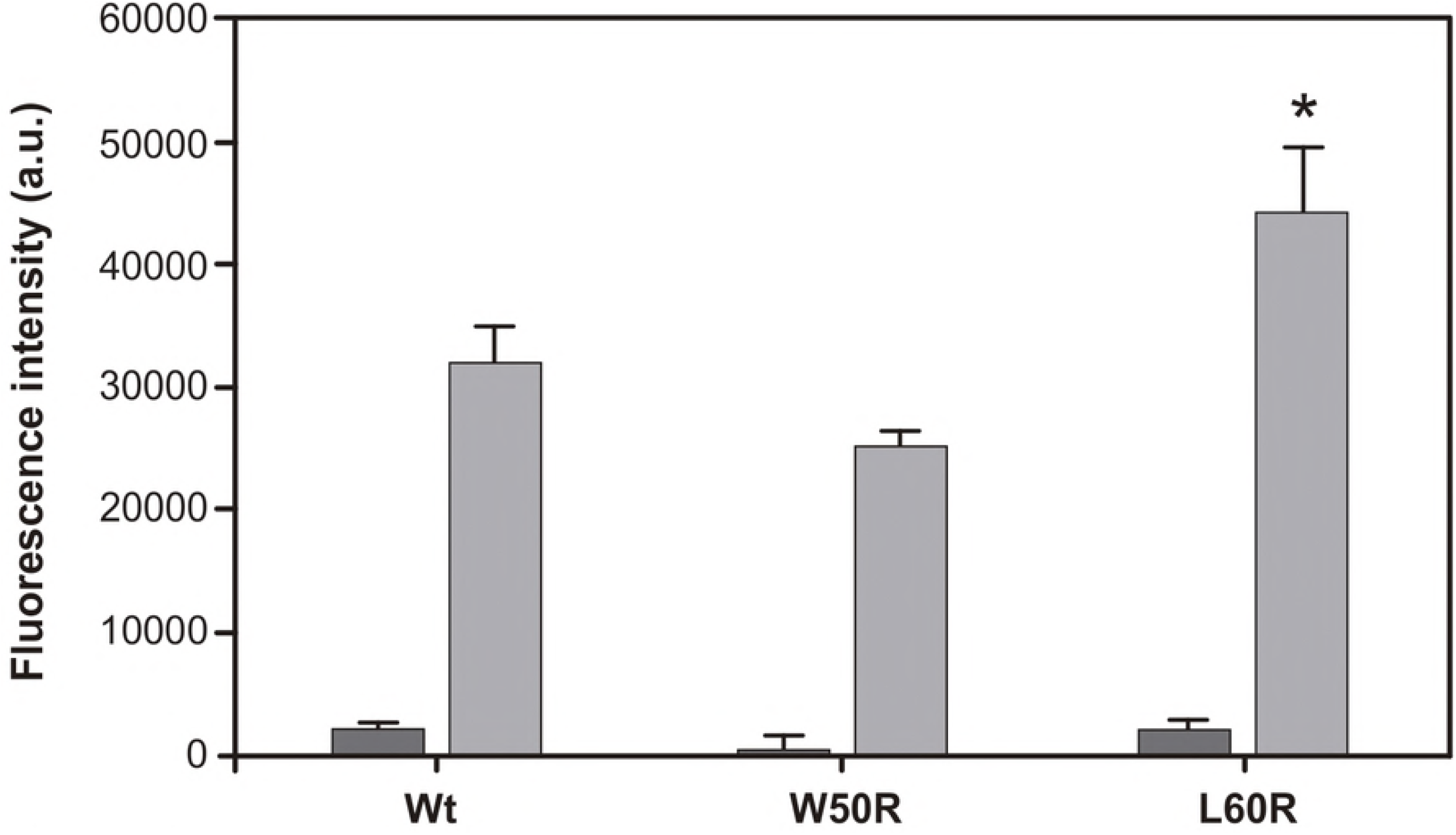
Protein binding to SDS. ApoA-I were incubated at a concentration of 0.2 mg/mL in Citrate phosphate buffer pH 7.4 in the absence (dark gray bars) or in the presence (clear gray bars) of 0.2 mM SDS. After 48 h at 37°C ThT was added to a 1:1 molar ratio to protein and relative fluorescence quantified as described in Figure 5D. The symbol * denotes difference with respect to Wt at P<0.05.

### Effect of mutations on lipid clearance and cellular responses

It is clear that the conformational flexibility could alter the equilibrium between function and toxicity, and thus we set out to test whether the structural shift detected in the N terminus of L60R could affect the protein interaction with a model of membrane. It is well known that the analysis of MLV clearance at the lipid transition temperature is a traditional functional parameter to test efficiency of apoA-I for lipids solubilization. Fig 7 shows that the L60R mutant clears DMPC MLV with a small by significantly higher efficiency than the Wt indicating a conserved function in lipid binding.

**Fig 7.**
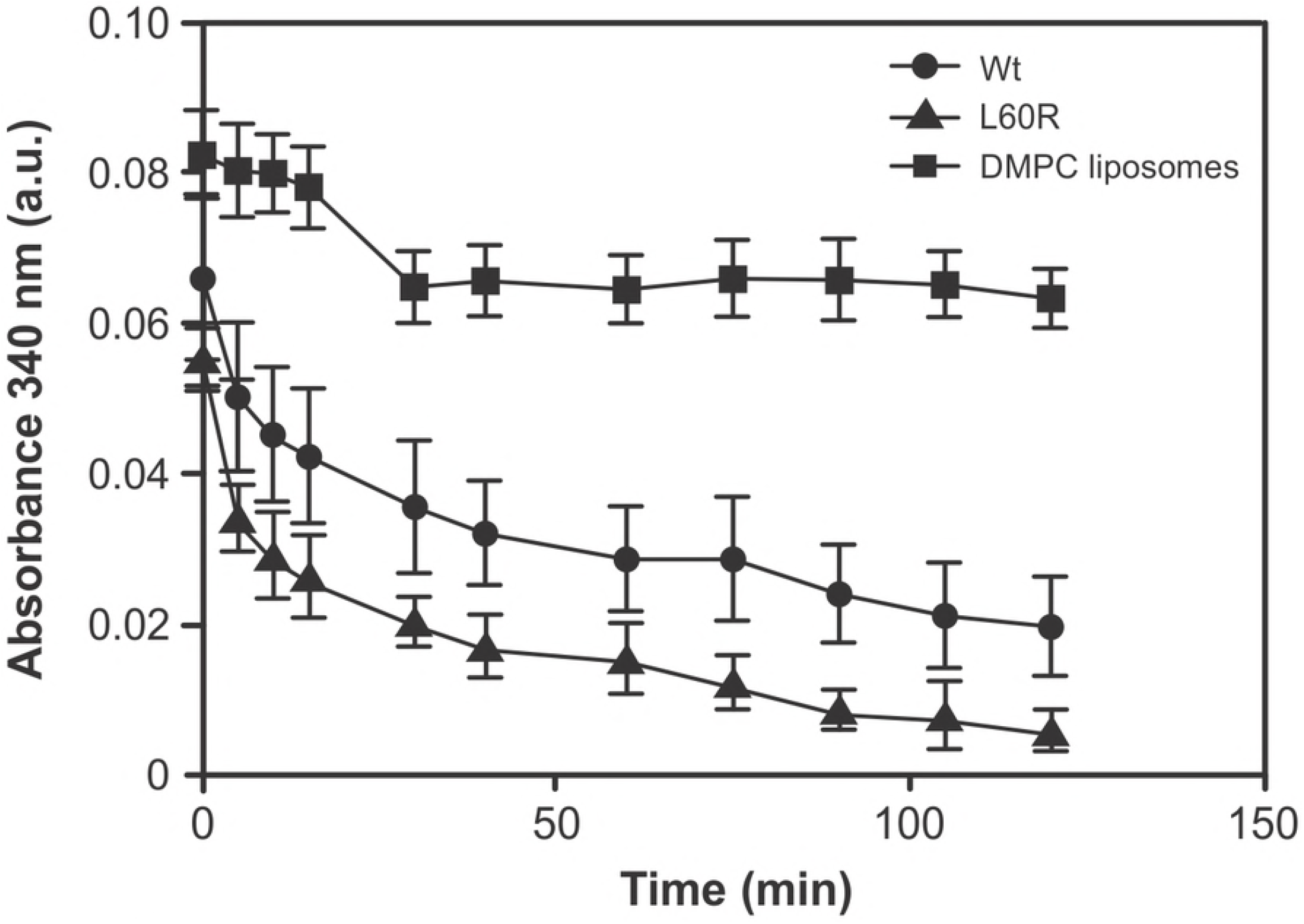
Characterization of the mutant L60R on the spontaneous formation of lipid:protein complexes. Multilamellar DMPC liposomes were incubated at 24°C in the presence of Wt (circles) or L60R (triangles) at a 80:1 lipid to protein molar ratio. Absorbance was measured at 340 nm in the Microplate Reader. Significant difference among both variants was determined by comparing absorbance at the last point of the measured kinetics (120 min). L60R Absorbance was different with respect to Wt at P<0.05 as measured by the Student’s Test.

Finally, and in order to test the hypothesis that apoA-I variants could induce cellular responses associated to a pro-inflammatory landscape, we compared the release of TNF-α and IL-1β from THP-1 cells, which is widely used as a human model of macrophages. As Fig 8A and B show, while W50R behaves as the Wt, L60R induced the release of these cytokines. MTT reduction measurements indicated that cell viability was preserved under the different conditions tested (not shown)

**Fig 8.**
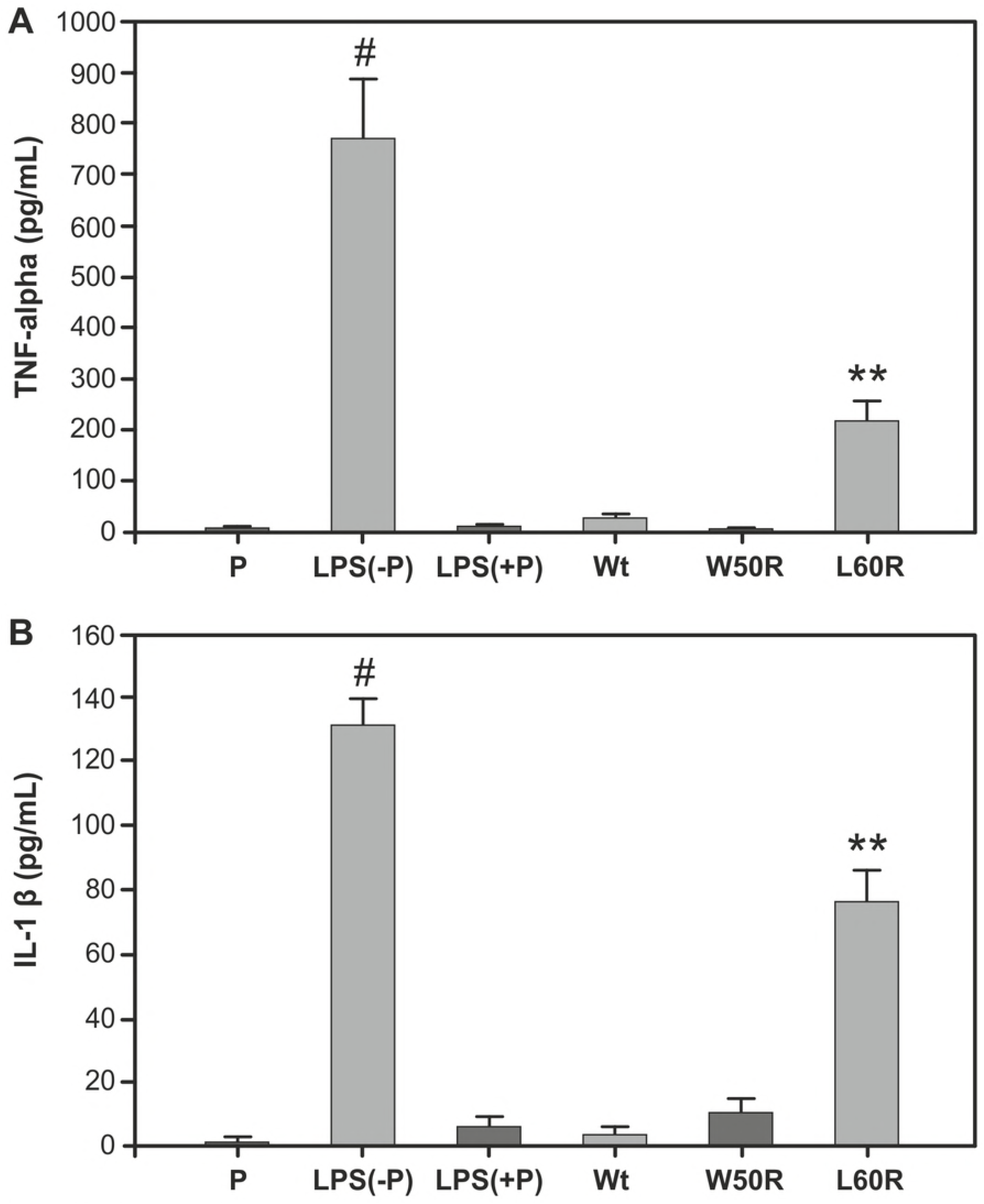
Induction of TNF-α (A) and IL-1β (B) release from cultured macrophages. THP-1 human monocytes were activated to macrophages by the addition of 5 ng/mL of Phorbol esters for 48 h. One μg/mL (A) or 0.5 μg/mL (B) of Wt, W50 or L60R were incubated with the cells for 3 h in the presence of Polymyxin B. Incubation of cells with LPS, either in the presence (LPS +P) or absence (LPS-P) of Polymyxin B, was used as negative and positive control respectively. P represents an extra negative control in which cells are incubated only in the presence of Polymyxin B. Symbol #represents significant difference respect to negative control (LPS +P) at P< 0.05. Symbol ** represents significant difference with respect to Wt at P< 0.001.

## Discussion

Structural flexibility is essential for apolipoproteins in order to fulfill complex functions, as it is required for apoA-I to interact within micro environmental ligands to solubilize lipids; nevertheless, this property may put the proteins into the risk of suffering subtle structural shifts from the native structure. The late-onset of the clinical manifestations of the amyloidosis disease due to apoA-I variants supports the fact that the final fibrillar conformation detected in the lesions is probably the result of progressive events that cooperate to give rise to the insoluble protein aggregates. Thus, it could be possible that a higher yield of partially folded variants could work late as seeding events in the long term period.

The structural modifications that result from the natural mutations in apoA-I are not easily predicted. Interestingly, although G26R, W50R, and L60R experiment a similar substitution in the N terminus (the replacement of a neutral amino acid by an Arg), only little disorder is observed for G26R [16] and W50R (this work and others ([16]; instead the structural arrangement observed herein for L60R is more evident. Das et al suggested that more than a strong conformational shift, the mutations in apoA-I may induce perturbations, increasing the accessibility of amyloid-prone segments that favor protein aggregation [16]. Such mild perturbations could increase the cleft from the α helical bundles allowing the accessibility to not-yet-described proteases which help the release of the peptides identified in the lesions. By an elegant model, they propose that desestabilization of the four-helix bundle containing residues may favor the formation of a resistant amyloid core by interactions of the N-terminal amyloid hot spots.

In agreement with Das et al. [16], our fluorescence and dichroism spectroscopic measurements (a small red shift in the Trp environment and a preserved secondary structure) suggest a mild effect of the Trp substitution by an Arg in position 50.Mutation W50R is comprised in the middle of the segment L44-S55 which was shown in the crystal structure having an extended conformation consistent with the β-strand-like geometry (Fig 9). It was proposed that the exposure of this segment could initiate the α-helix to β-sheet apoA-I conversion in amyloidosis. Nevertheless, the disruptive presence of a positive charge in replacement of the aromatic Trp is detected in our experimental design by an increased yield in the Bis-ANS binding and minor change in the near UV spectra, together with the detection of stronger binding to heparin at acidic pH. It is worth mentioning that the N-terminal 1–83 fragment of W50R variant was shown to participate in heparin-mediated fibril formation [39].

**Fig 9.**
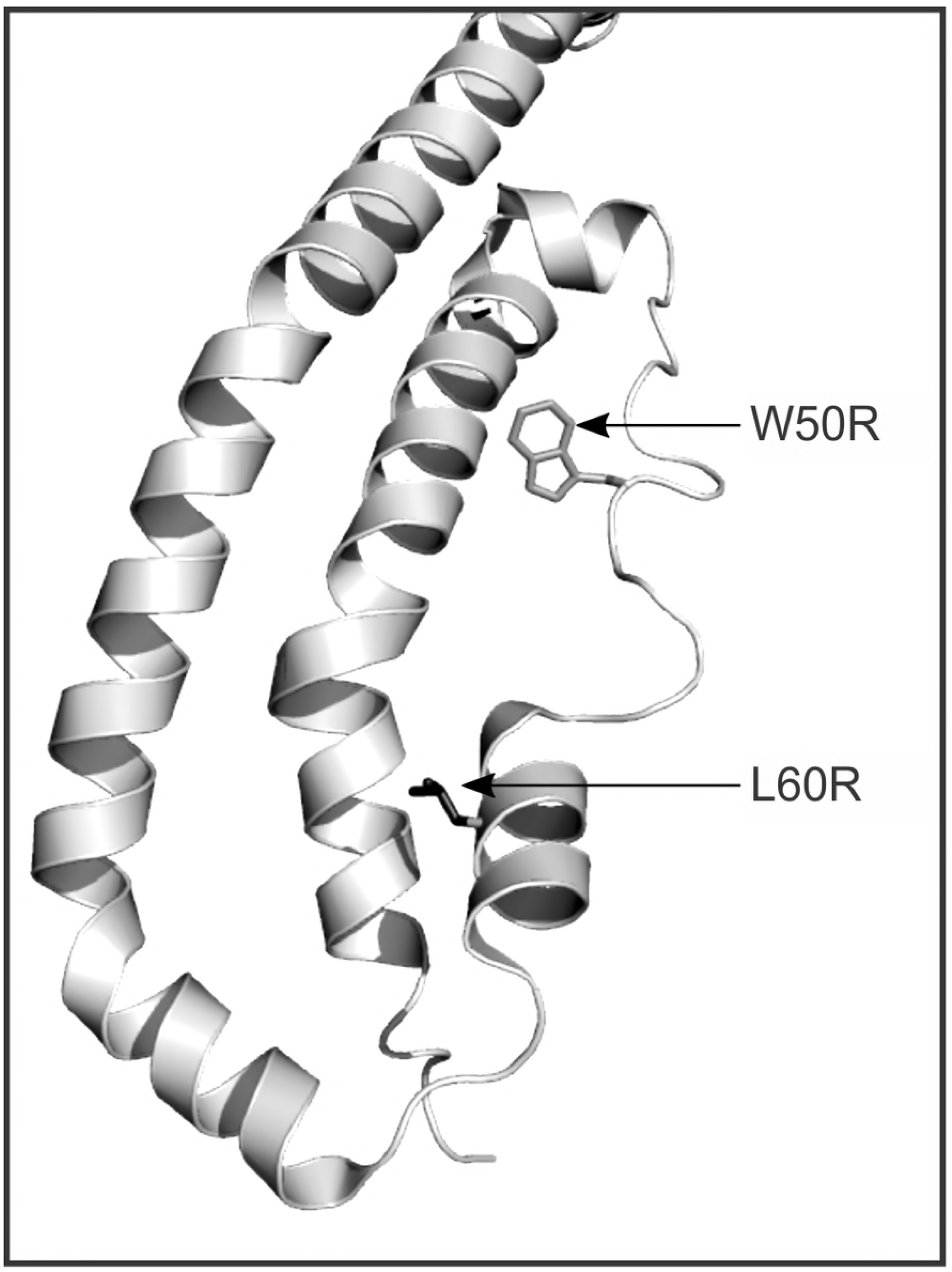
Locations of amyloidogenic mutations W50R and L60R in the structure of apoA-I. The structure was obtained by the PyMOL Molecular Graphics System, Version 2.0 (Schrödinger, LLC), from the X-ray crystal structure of Δ(185-243) apoA-I (PDB ID 3R2P). N terminal residues subjected to mutations are shown in stick representation.

Acidic intracellular milieu was associated to macrophages activation within inflammatory lesions [40]. As mentioned above, the W50R variant is one of the few exceptions in which apoA-I mutants’ deposits are associated to the glomeruli [13][41]. The extracellular matrix (ECM), especially the proteoglycans (PGs) have diverse biologic functions, including binding of growth factors and regulation of collagen fibrillogenesis [42], and their composition is tissue specific [43]. In addition, it was found that decorin (a dermatan/chondroitin sulfate PG) accumulated in amyloid deposits, but not in deposits of fibrillary glomerulo nephritis [44]. From our previous studies we learnt that binding of proteins to GAGs seems to be a cooperative, specific interaction [45]. We have previously observed that the amyloidogenic mutant R173P shows sensitive binding to heparin under physiological pH in spite of the loss in one positive charge, probably due to the exposition of cryptic positive residues which are accessible by the break in the amphipatic helix induced by the Proline residue in position 173 [15]. Even though binding at physiologic pH is not clear for the mutants tested here it is expected that the acidic milieu may strength protein-GAGs interactions. Although further studies are worth to be done, it could be speculated that the positive Arg charge could induce a stronger retention within the glomeruli’s GAGs thus helping its aggregation, especially under situations of kidney disease associated to inflammation.

As mentioned above, L60R shows (as compared to Wt), a red shift in Trp fluorescence which may indicate a relatively higher exposure of these aromatic residues to the polar environment. Due to the average contribution of the 4 Trp in the native structure, this shift may indicate either a subtle movement of side chains or larger scale conformational changes of the protein. The observed lower binding to Bis-ANS may suggest that the disruption of the bottom hydrophobic cluster by the positive charge of the Arg (Fig 9) could bring this amino acid more exposed, as it results from a higher binding to SDS which is not detected for the other amyloidogenic mutant tested in this work (Fig 6); moreover, this structural disorder should allow the permeation of proteases, as it is shown here. Although trypsin is not a physiological protease of apoA-I in circulation, it helps to get insight into protein structure. In order to compare trypsin induced proteolysis, we analyzed by the expassy software (www.expasy.org) the predicted sites to be substrate of this enzyme. Arg in position 60 should separate only one amino acid from a predicted fragment of three residues (59-61). Nevertheless, a higher efficiency of this variant’s processing is observed (Fig 2); the same tendency is detected under this experimental design by using MMP-12. Altogether, the rearrangement of the Trp environment in the L60R mutant, the higher accessibility to proteases and the lower CD spectral signal at 255-280 nm respect to the Wt agree with a spatial rearrangement with a decreased stability and an increase in protein flexibility. The increased susceptibility to proteases could explain the appearance of a 10 kDa molecular mass N terminal peptide within the lesions [8], although interestingly *in vitro* studies with the 93-residue N-terminal fragment demonstrated that the peptide freed behaves similar to Wt in the aggregation pattern than the same peptide with the Wt sequence [46].

In addition to protein misfolding, it is worth to consider the possibility that at least part of the clinical manifestations could be due to a loss in protein function, such as lipid binding, which in addition could increase the amount of lipid-free protein, amyloid-prone precursors. In order to consider this possibility, we first modeled the domain in which the mutation occurs (amino acids 52-65) in a helix wheel modeling software (http://lbqp.unb.br/NetWheels/). The modeling predicts that due to the periodicity that brings in close proximity residues Val53, Leu64 and Phe57, the Leu 60 contributes to form the apolar phase of the putative amphipatic class A α-helix (Fig 10).

**Fig 10.**
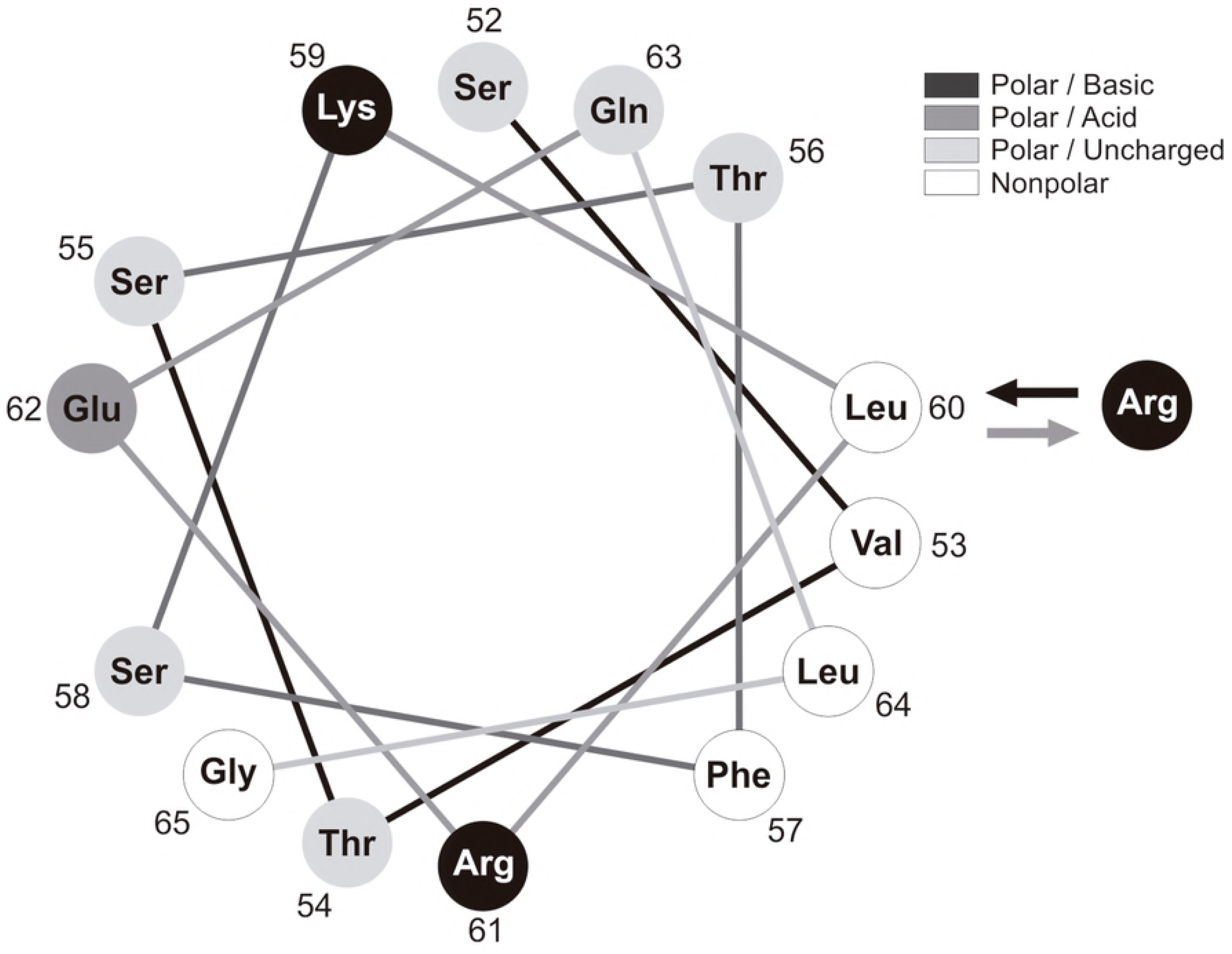
Helical wheel model of the putative α-helix comprising residues 52-65 of apoA-I. Wt sequence represented as an helix seen down the long axis, with an amino acid arrangement considering an ideal α-helix (l00^°^ rotation per amino acid). Gray scale in the Figure corresponds to the nature of different amino acids. Toward the right side of the helix it is indicated the replacement of Arg in position 60 in the nonpolar phase.

The positive charge of the Arg may thus perturb the hydrophobic bottom cluster. The mutation in position 60 should in addition disrupt a Leu zipper stabilized with other Leu residues in the same molecule or with a second molecule in the native dimer [47]. In our hands the shift among dimer and monomer conformation was not dramatically modified (S. Fig 1). Moreover, this effect is not drastic enough to decrease the efficiency to solubilize neutral lipids (Fig 7) as it is the case with W50R and G26R substitutions [16]. Instead, it could be that the relative exposition of the positive charge in this segment could favor its interactions and/or shift toward a pathological amyloidlike structure when interacting with other ligands, as negative lipids, as it suggested by the SDS binding experiments.

Finally, we have recently shown that other N terminal apoA-I mutant (Gly26Arg), induces the release of TNF-α and IL-1β from a model of macrophages [14], in a pathway probably involving the specific activation of the NF-κB pro-inflammatory cascade [48]. The kidney is a major target organ of innate immune inflammatory diseases. The deposition of the acute-phase reactant serum amyloid A (SAA) as amyloid causes progressive glomerular and vascular damage and leads to organ failure [49]. It was suggested that this and other misfolded proteins could be recognized by pattern recognition receptors (PRRs) resulting in the activation of pro inflammatory cascades, being increased IL-1β secretion responsible for most of the systemic features of this group of disorders [50]. IL-1β production may induce the synthesis of other cytokines as TNF-α. Our results shown here indicate that mild conformational rearrangement detected for L60R variant may induce the activation of the microenvironment toward a pro-inflammatory landscape which could help to perpetuate events triggering organ damage.

In conclusion, in agreement with other groups, our data support that it is not required a large overall destabilization of the tertiary structure of apoA-I to become amyloidogenic; either reduced protection of the major amyloid “hot spot”, increased susceptibilities to proteases or partial oxidation could induce cooperative shift toward a misfolded conformation. Nevertheless, the chronic clinical phenotype indicates that the landscape in which protein circulates (molecular crowding, acidification, oxidation, interactions with ligands etc) could play key roles to give rise to amyloid species. These species could either be nucleus of aggregation or, as we have previously suggested, work as signal receptors starting up cellular events associated to the pathology. Further research will help to explore different pathways.

## Acknowledgements

Authors acknowledge Mr. Mario Ramos for invaluable help with Figures’ design. We include acknowledgements to Mrs Rosana del Cid for English assistance, Lic Letizia Bauzá for FPLC measurements, to Lic Tomás Masson for help with Mass Spec analysis and Miss Ana L. Rodriguez for technical assistance.

## References

1. Schaefer EJ, Anthanont P, Asztalos BF. High-density lipoprotein metabolism, composition, function, and deficiency. Curr Opin Lipidol. England; 2014;25: 194–199. doi:10.1097/MOL.0000000000000074

2. Wu BJ, Ong KL, Shrestha S, Chen K, Tabet F, Barter PJ, et al. Inhibition of arthritis in the lewis rat by apolipoprotein A-I and reconstituted high-density lipoproteins. Arterioscler Thromb Vasc Biol. 2014;34: 543–551. doi:10.1161/ATVBAHA.113.302832

3. Barter PJ, Nicholls S, Rye KA, Anantharamaiah GM, Navab M, Fogelman AM. Antiinflammatory properties of HDL. Circ Res. 2004;95: 764–772. doi:10.1161/01.RES.0000146094.59640.13

4. Assmann G, Schmitz G, Funke H, Von Eckardstein A. Apolipoprotein A-I and HDL deficiency. Curr Opin Lipidol. 1990;1: 110–115. doi:10.1097/00041433-199004000-00005

5. Sorci-Thomas MG, Thomas MJ. The effects of altered apolipoprotein A-I structure on plasma HDL concentration. Trends Cardiovasc Med. United States; 2002;12: 121–128.

6. Van Allen MW, Frohlich JA, Davis JR. Inherited predisposition to generalized amyloidosis. Clinical and pathological study of a family with neuropathy, nephropathy, and peptic ulcer. Neurology. United States; 1969;19: 10–25.

7. Booth DR, Tan SY, Booth SE, Hsuan JJ, Totty NF, Nguyen O, et al. A new apolipoprotein Al variant, Trp50Arg, causes hereditary amyloidosis. QJM. England; 1995;88: 695–702.

8. Soutar AK, Hawkins PN, Vigushin DM, Tennent GA, Booth SE, Hutton T, et al. Apolipoprotein AI mutation Arg-60 causes autosomal dominant amyloidosis. Proc Natl Acad Sci U S A. United States; 1992;89: 7389–7393.

9. Rowczenio D, Stensland M, de Souza GA, Strøm EH, Gilbertson JA, Taylor G, et al. Renal Amyloidosis Associated With 5 Novel Variants in the Fibrinogen A Alpha Chain Protein. Kidney Int Reports. 2017;2: 461–469. doi:10.1016/j.ekir.2016.11.005

10. Kidd J, Carl DE. Renal amyloidosis. Current Problems in Cancer. 2016. pp. 209–219. doi:10.1016/j.currproblcancer.2016.08.002

11. Gregorini G, Izzi C, Obici L, Tardanico R, Röcken C, Viola BF, et al. Renal apolipoprotein A-I amyloidosis: a rare and usually ignored cause of hereditary tubulointerstitial nephritis. J Am Soc Nephrol. 2005;16: 3680–6. doi:10.1681/ASN.2005040382

12. Gregorini G, Izzi C, Ravani P, Obici L, Dallera N, Del Barba A, et al. Tubulointerstitial nephritis is a dominant feature of hereditary apolipoprotein A-I amyloidosis. Kidney Int. 2015;87: 1223–1229. doi:10.1038/ki.2014.389

13. Tougaard BG, Pedersen KV, Krag SR, Gilbertson J a, Rowczenio D, Gillmore JD, et al. A case report of hereditary apolipoprotein A-I amyloidosis associated with a novel APOA1 mutation and variable phenotype. Eur J Med Genet. 2016;59: 474–7. doi:10.1016/j.ejmg.2016.05.015

14. Ramella Na, Schinella GR, Ferreira ST, Prieto ED, Vela ME, Ríos JL, et al. Human apolipoprotein A-I natural variants: molecular mechanisms underlying amyloidogenic propensity. PLoS One. 2012;7: e43755. doi:10.1371/journal.pone.0043755

15. Rosú SA, Rimoldi OJ, Prieto ED, Curto LM, Delfino JM, Ramella NA, et al. Amyloidogenic propensity of a natural variant of human apolipoprotein A-I: Stability and interaction with ligands. PLoS One. 2015;10: 1–17. doi:10.1371/journal.pone.0124946

16. Das M, Mei X, Jayaraman S, Atkinson D, Gursky O. Amyloidogenic mutations in human apolipoprotein A-I are not necessarily destabilizing - a common mechanism of apolipoprotein A-I misfolding in familial amyloidosis and atherosclerosis. FEBS J. England; 2014;281: 2525–2542. doi:10.1111/febs.12809

17. Das M, Wilson CJ, Mei X, Wales TE, Engen JR, Gursky O. Structural Stability and Local Dynamics in Disease-Causing Mutants of Human Apolipoprotein A-I: What Makes the Protein Amyloidogenic? J Mol Biol. 2016;428: 449–462. doi:10.1016/j.jmb.2015.10.029

18. Tricerri MA, Behling Agree AK, Sanchez SA, Bronski J, Jonas A. Arrangement of apolipoprotein A-I in reconstituted high-density lipoprotein disks: an alternative model based on fluorescence resonance energy transfer experiments. Biochemistry. 2001;40: 5065–5074. doi:bi002815q [pii]

19. Ramella Na, Rimoldi OJ, Prieto ED, Schinella GR, Sanchez S a, Jaureguiberry MS, et al. Human apolipoprotein A-I-derived amyloid: its association with atherosclerosis. PLoS One. 2011;6: e22532. doi:10.1371/journal.pone.0022532

20. Prieto ED, Ramella N, Cuellar LA, Tricerri MA, Garda HA. Characterization of a human apolipoprotein a-I construct expressed in a bacterial system. Protein J. Netherlands; 2012;31: 681–688. doi:10.1007/s10930-012-9448-z

21. Gomori G. Preparation of Buffers for Use in Enzyme Studies (by G. Gomori). Enzyme. 2004. pp. 2–10. doi:10.1016/0076-6879(55)01020-3

22. Lakowicz JR. Principles of Fluorescence Spectroscopy. Springer New York; 2006.

23. Schmid FX. Spectral methods of characterizing protein conformation and conformational changes. Protein structure A practical approach. 1989. pp. 251–285.

24. LeVine H. Quantification of β-sheet amyloid fibril structures with thioflavin T. Methods Enzymol. 1999;309: 274–284. doi:10.1016/S0076-6879(99)09020-5

25. Lindgren M, Sörgjerd K, Hammarström P. Detection and characterization of aggregates, prefibrillar amyloidogenic oligomers, and protofibrils using fluorescence spectroscopy. Biophys J. 2005;88: 4200–4212. doi:10.1529/biophysj.104.049700

26. Wetzel R, Chemuru S, Misra P, Kodali R, Mukherjee S, Kar K. An aggregate weight-normalized thioflavin-t measurement scale for characterizing polymorphic amyloids and assembly intermediates. Methods in Molecular Biology. 2018. pp. 121–144. doi:10.1007/978-1-4939-7811-3_6

27. Ellis RA. Principles and techniques of electron microscopy: Biological applications, Volume 9. Cell. 1979;17: 235–236. doi:10.1016/0092-8674(79)90312-X

28. Tulumello D V., Deber CM. SDS micelles as a membrane-mimetic environment for transmembrane segments. Biochemistry. 2009;48: 12096–12103. doi:10.1021/bi9013819

29. Park EK, Jung HS, Yang HI, Yoo MC, Kim C, Kim KS. Optimized THP-1 differentiation is required for the detection of responses to weak stimuli. Inflamm Res. 2007;56: 45–50. doi:10.1007/s00011-007-6115-5

30. Bradford MM. A rapid and sensitive method for the quantitation of microgram quantities utilizing the principle of …. Anal Biochem. 1976;72: 248–254.

31. Davidson WS, Arnvig-McGuire K, Kennedy A, Kosman J, Hazlett TL, Jonas A. Structural organization of the N-terminal domain of apolipoprotein A-I: Studies of tryptophan mutants. Biochemistry. 1999;38: 14387–14395. doi:10.1021/bi991428h

32. Martins SM, Chapeaurouge A, Ferreira ST. Folding intermediates of the prion protein stabilized by hydrostatic pressure and low temperature. J Biol Chem. United States; 2003;278: 50449–50455. doi:10.1074/jbc.M307354200

33. Mei X, Atkinson D. Crystal Structure of C-terminal Truncated Apolipoprotein A-I Reveals the Assembly of High Density Lipoprotein (HDL) by. 2011. doi:10.1074/jbc.M 111.260422

34. Sreerama N, Woody RW. Estimation of protein secondary structure from circular dichroism spectra: Comparison of CONTIN, SELCON, and CDSSTR methods with an expanded reference set. Anal Biochem. 2000;287: 252–260. doi:10.1006/abio.2000.4880

35. Cohlberg JA, Li J, Uversky VN, Fink AL. Heparin and other glycosaminoglycans stimulate the formation of amyloid fibrils from ??-synuclein in vitro. Biochemistry. 2002;41: 1502–1511. doi:10.1021/bi011711s

36. Aguilera JJ, Zhang F, Beaudet JM, Linhardt RJ, Colón W. Divergent effect of glycosaminoglycans on the in vitro aggregation of serum amyloid A. Biochimie. Elsevier; 2014;104: 70–80.

37. Oberkersch R, Attorresi AI, Calabrese GC. Low-molecular-weight heparin inhibition in classical complement activaton pathway during pregnancy. Thromb Res. 2010;125. doi:10.1016/j.thromres.2009.11.030

38. Ahmad MF, Ramakrishna T, Raman B, Rao CM. Fibrillogenic and Non-fibrillogenic Ensembles of SDS-bound Human α-Synuclein. J Mol Biol. 2006;364: 1061–1072. doi:10.1016/j.jmb.2006.09.085

39. Mikawa S, Mizuguchi C, Nishitsuji K, Baba T, Shigenaga A, Shimanouchi T, et al. Heparin promotes fibril formation by the N-terminal fragment of amyloidogenic apolipoprotein A-I. FEBS Lett. 2016;590: 3492–3500. doi:10.1002/1873-3468.12426

40. Oorni K, Rajamaki K, Nguyen SD, Lahdesmaki K, Plihtari R, Lee-Rueckert M, et al. Acidification of the intimal fluid: the perfect storm for atherogenesis. J Lipid Res. 2015;56: 203–214. doi:10.1194/jlr.R050252

41. Samillán-sosa R, Sención-martínez G, Lopes-martín V, Martínez-gonzález MA, Solé M, Luis J, et al. Amiloidosis renal hereditaria por depósito de apolipoproteína AI: un reto diagnóstico. NEFROLOGíA. Sociedad Española de Nefrología; 2015;35: 322–327. doi:10.1016/j.nefro.2015.05.002

42. Iozzo R V. MATRIX PROTEOGLYCANS: From Molecular Design to Cellular Function. Annu Rev Biochem. 1998;67: 609–652. doi:10.1146/annurev.biochem.67.1.609

43. Schaefer L, Gröne HJ, Raslik I, Robenek H, Ugorcakova J, Budny S, et al. Small proteoglycans of normal adult human kidney: Distinct expression patterns of decorin, biglycan, fibromodulin, and lumican. Kidney Int. 2000;58: 1557–1568. doi:10.1046/j.1523-1755.2000.00317.x

44. Stokes MB, Holler S, Cui Y, Hudkins KL, Eitner F, Fogo A, et al. Expression of decorin, biglycan, and collagen type I in human renal fibrosing disease. Kidney Int. 2000;57: 487–498. doi:10.1046/j.1523-1755.2000.00868.x

45. Rosú SA, Toledo L, Urbano BF, Sanchez SA, Calabrese GC, Tricerri MA. Learning from Synthetic Models of Extracellular Matrix; Differential Binding of Wild Type and Amyloidogenic Human Apolipoprotein A-I to Hydrogels Formed from Molecules Having Charges Similar to Those Found in Natural GAGs. Protein J. 2017;36. doi:10.1007/s10930-017-9728-8

46. Raimondi S, Guglielmi F, Giorgetti S, Di Gaetano S, Arciello A, Monti DM, et al. Effects of the known pathogenic mutations on the aggregation pathway of the amyloidogenic peptide of apolipoprotein A-I. J Mol Biol. England; 2011;407: 465–476. doi:10.1016/j.jmb.2011.01.044

47. Gursky O, Mei X, Atkinson D. The Crystal Structure of the C-Terminal Truncated Apolipoprotein A-I Sheds New Light on Amyloid Formation by the N-Terminal Fragment. 2012;

48. Ramella NA, Andújar I, Ríos JL, Rosú SA, Tricerri MA, Schinella GR. Human apolipoprotein A-I Gly26Arg stimulation of inflammatory responses via NF-kB activation: Potential roles in amyloidosis? Pathophysiology. 2018; doi:10.1016/j.pathophys.2018.08.002

49. Scarpioni R, Obici L. Renal involvement in autoinflammatory diseases and inflammasome-mediated chronic kidney damage. Clin Exp Rheumatol. 2018;36: S54–S60.

50. Gustot A, Raussens V, Dehousse M, Dumoulin M, Bryant CE, Ruysschaert JM, et al. Activation of innate immunity by lysozyme fibrils is critically dependent on cross-ß sheet structure. Cell Mol Life Sci. 2013;70: 2999–3012. doi:10.1007/s00018-012-1245-5

